# Persistent effects of repeated adolescent and adult heroin vapor inhalation in female Wistar rats

**DOI:** 10.1101/2024.05.06.592492

**Authors:** Arnold Gutierrez, Michael A. Taffe

**Affiliations:** Department of Psychiatry, University of California, San Diego; La Jolla, CA, USA

## Abstract

Adolescent drug exposure has been associated with more severe mental health outcomes related to substance abuse and anxiety disorders. The aim of the present study was to contrast the long-term effects of repeated heroin vapor inhalation during adolescence with similar heroin exposure in adulthood. Groups of female Wistar rats underwent twice daily 30-minute sessions of heroin or propylene glycol (control) vapor inhalation from postnatal days (PND) 36-45 or PND 85-94, respectively. Nociception was assessed after vapor inhalation sessions and forty days later, for the Adolescent-Exposed and Adult-Exposed groups. Anxiety-like behavior was assessed with an elevated plus-maze (EPM) and spatial learning was assessed with a Barnes maze. Acute effects of naloxone (0.3 mg/kg, i.p.) and heroin (0.5 and 1.0 mg/kg, s.c.) on thermal nociception were determined on PND 140/189 and PND 149/198, respectively. Repeated heroin vapor inhalation produced anti-nociceptive tolerance across sessions in both adolescent and adult rats, with the adolescents exhibiting more complete tolerance. Heroin vapor inhalation produced anxiolytic effects, regardless of age of exposure. There were no effects of heroin on spatial learning. Naloxone produced acute hyperalgesia in all but the Adolescent-Exposed heroin group, and heroin anti-nociception was blunted in both heroin-exposed groups at the highest heroin dose. Repeated heroin vapor inhalation can produce lasting effects on nociception and anxiety-like behavior that persist for months after the exposure. Importantly, these findings suggest that adolescent exposure to heroin vapor produces specific effects on nociception that are not observed when exposure occurs in adulthood.

## INTRODUCTION

Inhalation is a route of drug administration common to many that use opioid drugs nonmedically (Parent et al. 2021). We and others have reported success in using Electronic Nicotine Delivery Systems (ENDS), or more generally termed Electronic Drug Delivery Systems (EDDS), to deliver behaviorally active doses of opioids by inhalation in preclinical rodent models (Gutierrez et al. 2021; Gutierrez et al. 2022b; Gutierrez et al. 2020; Marchette et al. 2023; McConnell et al. 2021; Moussawi et al. 2020; Vendruscolo et al. 2018). EDDS technology has also been used to probe the lasting effects of repeated adolescent vapor inhalation of drugs such as nicotine (Gutierrez et al. 2024; Gutierrez et al. 2023), THC (Gutierrez et al. 2024; Nguyen et al. 2020), and heroin (Gutierrez et al. 2022a), and to determine lasting effects of *in utero* exposure to THC (Breit et al. 2022; Breit et al. 2020).

Adolescence is a critical neurodevelopmental period that is associated with heightened risk-taking behavior, including experimentation with intoxicating substances (Miech et al. 2023; Temourian et al. 2023), in humans; thus, adolescents are potentially at an increased risk for developing substance use disorders and experiencing drug-induced comorbidity with other affective disorders. For example, about 50% of US adult smokers started before the age of 18 and about 83% before the age of 21 (Ali et al. 2020). Also, problematic drug-taking in adolescence is associated with more severe problems with drugs in adulthood (McCabe et al. 2022), and adolescent drug use is associated with higher risk for developing anxiety disorders (Moylan et al. 2013). This is important because anxiety may itself drive drug-taking behavior, as it often precedes substance use problems (Bushnell et al. 2019) and is a predictor of relapse (Willinger et al. 2002). There are relatively few preclinical studies on the lasting effects of adolescent exposure to opioids on anxiety-like behavior, and results from the existing studies are mixed. For example, adolescent exposure to opioids has been reported to produce anxiolytic (Khani et al. 2022a), anxiogenic (Gutierrez et al. 2022a), or no anxiety-related effects (Lutz et al. 2013; Sanchez et al. 2016; Schwarz and Bilbo 2013) in adult rodents.

Associations have been reported between opioid abuse and cognitive dysfunction. For example, memory impairments resulting from opioid abuse have been reported in adult (Kroll et al. 2018; Ornstein et al. 2000) and adolescent humans (Vo et al. 2014). One recent study, which studied a number of cognitive domains in heroin users, found that cognitive impairment was specifically related to the route of administration) used (Ghosh et al. 2023); intravenous injection was associated with memory deficits while inhalation was associated with decreased ability to maintain attentional set. Injecting rodents with opioid drugs can likewise impair performance on tests of impulsivity (Pattij et al. 2009), cognitive flexibility (You et al. 2022), and working memory (Braida et al. 1994). Repeated opioid administration also degraded performance on tests of spatial navigation; however, the majority of such studies focus on behavioral outcomes of testing during ongoing treatment with repeated injections (Lu et al. 2010; Pu et al. 2002) or shortly after the discontinuation of a repeated opioid injection regimen during withdrawal (Luo et al. 2022). There is a dearth of data on the longer lasting effects of opioids on learning and memory in rodents, especially within the context of developmental exposure. One available recent study found that repeated injections of morphine during adolescence impaired measures of spatial learning and memory in the Morris water maze when male rats had reached adulthood (Khani et al. 2022b). It is presently unknown if these findings also extend to female rats or if opioid *inhalation* produces effects similar to those observed after injection. The importance of the latter is especially highlighted by the findings of Ghosh and colleagues (2023).

Opioid drugs are primarily used clinically for their analgesic effects. However, individuals that use opioids regularly often experience a decreased threshold for pain (Compton et al. 2001), even after drug cessation (Ho and Dole 1979). This hyperalgesic effect is associated with drug craving (Ren et al. 2009; Tsui et al. 2016) and may therefore increase the potential for relapse of non-medical users. Rodent studies have previously shown that adolescent opioid exposure can produce effects on opioid anti-nociception in early adulthood, including acceleration of tolerance across days of repeated morphine administration (Salmanzadeh et al. 2017) and blunted anti-nociception to oxycodone injection (Zhang et al.

2016). In a recent study, we confirmed that repeated *inhalation* exposure to heroin during adolescence alters baseline nociception and decreases anti-nociceptive effects of heroin that persist into middle adulthood in male and female rats (Gutierrez et al. 2022a). In that prior study, the effect of the age of the repeated heroin exposure was not investigated so any specific vulnerability of adolescents could not be determined. The major goal of the present study was therefore to investigate if there are differences in nociception, spatial learning, and anxiety-like behavior based on the age of heroin vapor inhalation exposure.

## METHODS

### Animals

Adolescent (N=16) and adult (N=16) female rats (Charles River) were used for this study. Rats arrived on Post-natal Day (PND) 29 or PND 78 and were habituated to the vivarium through PND 35 or PND 84, respectively, as previously described in (Gutierrez et al. 2022a). The vivarium was kept on a 12:12 hour light-dark cycle and behavior studies were conducted during the rats’ dark period. Food and water were provided ad libitum in the home cage, and body weights were recorded weekly beginning at PND 36 / PND 85. Experimental procedures were conducted in accordance with protocols approved by the IACUC of the University of California, San Diego and consistent with recommendations in the NIH Guide (Garber et al. 2011).

### Drugs

Heroin (diamorphine HCl) was dissolved in the propylene glycol (PG) vehicle for vapor inhalation or in physiological saline for subcutaneous (s.c.) injection. Naloxone (naloxone HCl) was dissolved in physiological saline and administered by intraperitoneal (i.p.) injection. For vapor inhalation sessions, a drug solution volume of ∼ 0.5 ml was loaded into the EDDS tanks. Fresh solutions were used for each inhalation session. Heroin was provided by the U.S. National Institute on Drug Abuse. Naloxone and propylene glycol were purchased from Fisher Scientific.

### Vapor drug treatment

Rats were exposed to vapor in sealed vapor exposure chambers (152 mm W X 178 mm H X 330 mm L; La Jolla Alcohol Research, Inc, La Jolla, CA, USA). E-vape controllers (Model SSV-3 or SVS-200; 58 watts; La Jolla Alcohol Research, Inc, La Jolla, CA, USA) were used to trigger Smok Baby Beast Brother TFV8 sub-ohm tanks equipped with V8 X-Baby M2 0.25 ohm coils. The exposure chambers were attached to vacuum exhaust lines that pulled ambient air through intake valves at ∼2 L per minute.

Rats underwent twice daily heroin (50 mg/mL in PG) or PG (control) vapor exposure sessions (30-minute duration) separated by a 6-hour inter-session interval beginning on PND 36 for the adolescents, or PND 85 for the adults. Vapor exposure sessions continued for total of 10 days (20 sessions; PND 36-45 or PND 85-94). Sessions consisted of six-second vapor deliveries triggered at five-minute intervals. Rats were exposed to vapor treatment in groups of four per chamber. The house vacuum line was turned on 30 seconds prior to vapor deliveries and closed immediately after. The total vapor clearance time was ∼5 minutes. Vapor began being cleared from the chamber 4.5 minutes after the last vapor delivery. This procedure was described previously described (Gutierrez et al. 2022a).

### Thermal nociception

#### Nociception assays

Tail-withdrawal tests were performed using a Branson Brainsonic CPXH Ultrasonic Bath (Danbury, CT). Water temperatures were set and maintained at 46°C, 48°C, 50°C, or 52°C (+/- 0.10°C) depending on the experiment. Prior to each measurement, the water bath was stirred, and the water temperature was verified by using a second thermometer. Withdrawal latencies were measured with a stopwatch, and a cutoff of 15 sec was used to avoid any potential for tissue damage.

#### Pre- and post-heroin inhalation nociception

Nociception assays were performed immediately before and after the first vapor inhalation session of the day on days 1 (session 1), 5 (session 9), and 10 (session 19). Thus, testing occurred on PND 36, 40, 45 for the Adolescent-Exposed rats and on PND 85, 89, 94 for the Adult-Exposed groups. The water temperature was maintained at 52°C for these assessments. The goals were to determine any age-related differences in the anti-nociceptive effects of heroin vapor exposure and to determine any age-related differences in the development of tolerance.

#### Baseline Nociception

Tail-withdrawal measurements were obtained at 12 weeks of age (PND 85-87) in groups treated as adolescents or at 19 weeks of age (PND 134-136) in groups treated as adults. Although all animals were adults by this point, to avoid confusion groups treated as adolescents shall be referred to as Adolescent-Exposed groups and those treated during adulthood as Adult-Exposed groups. A range of water temperatures (46°C-52°C) was used to assess baseline nociception, as previously described (Gutierrez et al. 2022a).Tail-withdrawal tests were performed over two days, with one day in between test days. Two temperatures were assessed per day, and measurements for each temperature were separated by 120 minutes. A counterbalanced order was used for the two temperature conditions for each test day.

Baseline nociception was again assessed in Adolescent and Adult groups at 17 weeks (PND 121) and 24 weeks (PND 170) of age, respectively. For this set of measurements, the target water temperature was 46°C. This temperature was selected based on the findings from our initial baseline nociception experiment and our previous work (Gutierrez et al. 2022a) showing that differences associated with repeated heroin vapor exposure are optimally detected at the lower range of temperatures.

#### Naloxone challenge

To assess the effects of opioid receptor antagonism on thermal nociception, rats were challenged with naloxone (0.3 mg/kg, i.p.) prior to warm water (46°C) tail-withdrawal measurements. Tail-withdrawal measurements were performed at 20 (PND 140) and 27 weeks (PND 189) of age in the Adolescent-Exposed and Adult-Exposed groups, respectively. Thermal nociception was measured immediately before naloxone (0.3 mg/kg, i.p.) was administered, and again 15 minutes post-injection.

#### Heroin challenge

Anti-nociceptive effects of heroin delivered via subcutaneous (s.c.) injection were assessed in the Adolescent-Exposed and Adult-Exposed rat groups at 21 (PND 149) and 28 (PND 198) weeks of age, respectively. The water bath was maintained at a temperature of 52°C for these experiments. Tail-withdrawal latencies were recorded immediately prior to heroin (0.5 mg/kg, s.c.) administration, and again 30 minutes after the first injection. A second heroin injection (1.0 mg/kg, s.c.) was administered 120 minutes after the first injection, and tail-withdrawal latencies were recorded 30 minutes after the second injection.

### Anxiety-like behavior

Anxiety-like behavior was assessed using an elevated plus-maze (EPM) test. Testing in the EPM was performed at 13 weeks (PND 93) and 20 weeks (PND 142) of age for the adolescent and adult groups, respectively. Testing was conducted in a dimly lit room. The apparatus consisted of two opposing closed arms and two opposing open arms perpendicular to the closed arms. The walls of the closed arms measured 40 cm in height. The apparatus was suspended 50 cm above the floor by four legs, each located toward the distal part of an arm. The test duration was five minutes. Behavior was recorded and measured using AnyMaze software (Stoelting, Wood Dale, IL). The EPM apparatus was cleaned with an ethanol solution after each subject completed testing.

### Spatial learning

#### Barnes maze apparatus

A Barnes maze apparatus consisting of a black circular platform (122 cm in diameter) was used to test spatial learning in rats. The apparatus had 20 holes, each measuring 10 cm in diameter, evenly spaced around the periphery. The top of the maze was 68.5 cm from the ground. Each hole, with the exception of the designated escape hole, was open, while the designated escape hole led to a removable black goal box.

#### Barnes maze procedure

Three days before commencing testing (PND 102 and 151 for the Adolescent- and Adult-Exposed groups, respectively), each rat was placed in an opaque container and transferred into the goal box through the escape hole. The container was used to minimize the animal’s exposure to the test room environment. The goal box was then covered, and each rat was allowed to habituate for a total of three minutes. The goal box was then removed with the rat inside, and the animal was transferred back to their home cage. The acquisition phase began three days after habituation and continued for five days (Adolescent-Exposed PND105-109; Adult-Exposed PND154-158). Each testing day consisted of two trials, with each trial totaling two minutes in duration. Testing was performed in cohorts of 5-6 rats at a time, and all rats in a group were required to complete the first trial prior to any rat beginning the second. The reversal phase (Adolescent-Exposed PND112-116; Adult-Exposed PND161-165) began two days after the completion of the last acquisition day. For the reversal phase, the location of the escape hole was rotated 180°. As with acquisition, reversal consisted of two trials per day, each trial totaling two minutes in duration. On each test day, trials commenced by transferring rats in a container to the center of the maze apparatus. Rats were confined to the container for 10 seconds before it was lifted, at which point the test began and behavior was measured and recorded using AnyMaze software (Stoelting, Wood Dale, IL). Upon completion of each trial, rats were returned to their cages to await the next trial or be returned to the vivarium, and the apparatus was thoroughly cleaned using an alcohol solution. For any rat that did not reach the escape location within the allotted time, a score equivalent to the maximum possible duration (i.e., 2 minutes) was given.

### Data and statistical analysis

Data were analyzed by Analysis of Variance (ANOVA) with repeated measures factors of Day for pre- and post-vapor inhalation session tail-withdrawal experiments and Barnes maze tests, Temperature in nociception experiments where multiple temperatures were assessed, Time in baseline nociception experiments that assessed measurements at different timepoints, and drug Treatment condition in the naloxone and heroin challenges. Mixed-effects analysis were utilized in circumstances where data were missing. Significant effects were followed by Sidak (two-level factors) or Tukey (multi-level factors) post hoc analyses. A criterion of P < 0.05 was used to infer significant differences. Analysis of the data collected on spatial learning in the Barnes maze included the first three days for each phase due to an apparent ceiling effect on learning by day three and a qualitative change in behavior observed that suggested increased exploratory behavior after the task had been mastered. Statistical analyses were performed using Prism 10.1.2 for Windows (GraphPad Software, Inc., San Diego, CA).

## RESULTS

### Thermal nociception

#### Pre-vapor nociception

The initial three-factor analysis of pre-session tail-withdrawal latencies confirmed effects of Age (F (1, 28) = 30.56, p < 0.0001) and Day (F (2, 56) = 7.04, p < 0.01); adults exhibited significantly longer latencies compared with adolescents, and latencies on Days 10 were significantly longer than on Day 1 (**Figure 1A**). There was also an interaction of Age with Day (F (2, 56) = 4.76, p < 0.05). The Tukey test confirmed that on Day 5, the adult rats exposed to heroin showed significantly longer latencies compared with the adolescent rats exposed to PG. By Day 10, the adult heroin-exposed rats showed significantly longer withdrawal latencies compared with each of the two adolescent groups. A follow-up two-way

**Figure 1.**
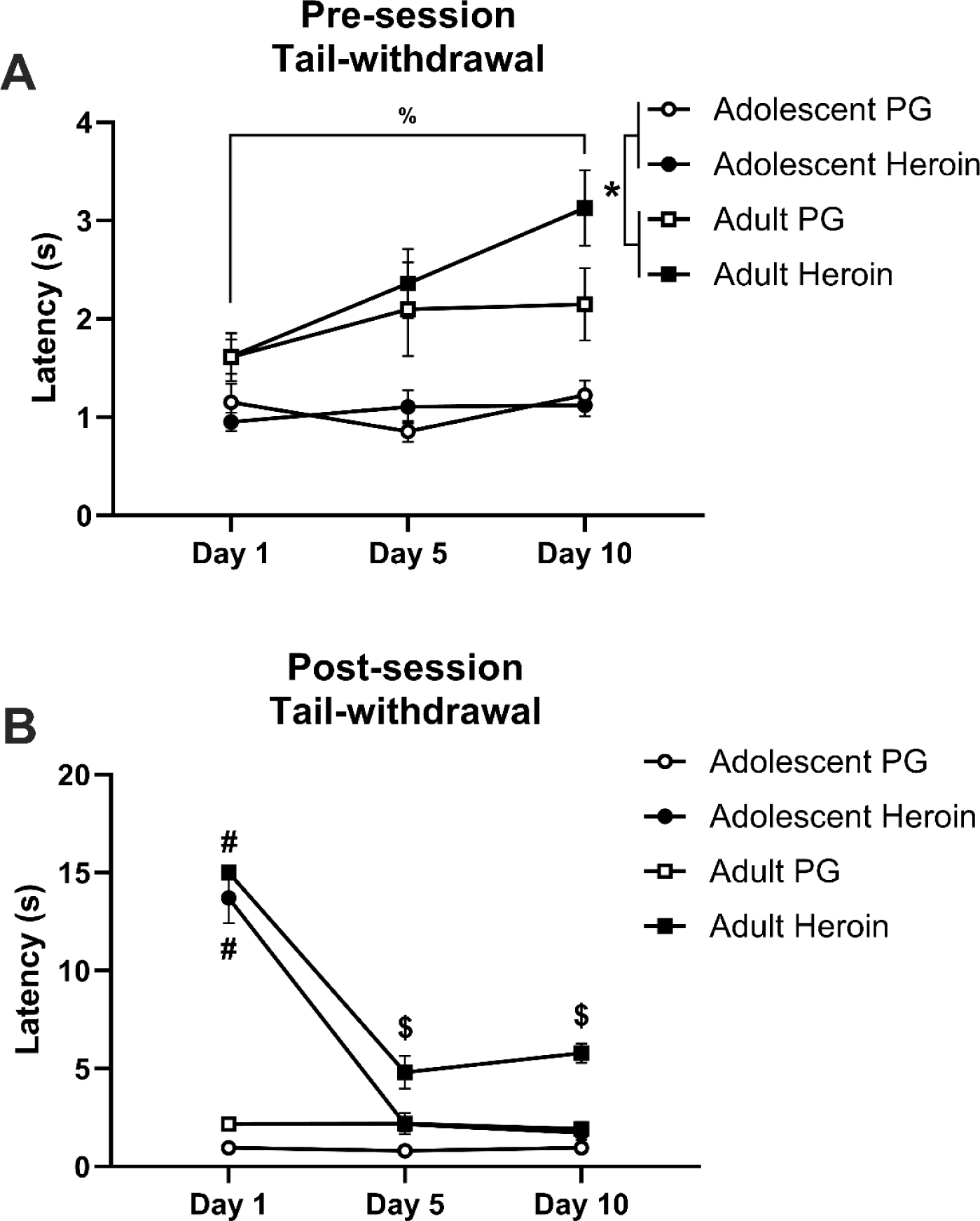
Mean (±SEM) tail-withdrawal latencies before and after the first vapor inhalation session of the day on days 1, 5, and 10 for adolescent (*N*=16; 8 PG, 8 Heroin) and adult (*N*=16; 8 PG, 8 Heroin) rats. Latencies recorded prior to (**A**) and after (**B**) inhalation sessions were measured at 52°C. A significant difference between the Adolescent and Adult groups is indicated with a *, a significant difference compared with all PG groups with #, a significant difference compared with all groups with $, and a significant difference between Days with %. analysis collapsed across Drug treatment again confirmed main effects of Age (F (1, 30) = 30.35, p < 0.0001) and Day (F (2, 60) = 6.74, p < 0.01), and the interaction of Age with Day (F (2, 60) = 4.56, p < 0.05). The post-hoc confirmed that the adolescents significantly differed from the adults on all three days and that Day 1 significantly differed from Days 5 and 10, only within the adults. No differences were confirmed within the adolescent groups.

#### Post-vapor nociception

Repeated heroin vapor exposure produced nociceptive tolerance differentially between adolescent and adult rats. The initial three-factor analysis of the post-session tail-withdrawal measurements confirmed main effects of Day (F (2, 56) = 156.43, p < 0.0001), Age (F (1, 28) = 32.92, p < 0.0001), and Drug (F (1, 28) = 285.74, p < 0.0001). The analysis also confirmed significant interactions of Drug with Day (F (2, 56) = 150.91, p < 0.0001) and Age (F (1, 28) = 4.77, p < 0.05). The Tukey test confirmed that on Day 1, each heroin group exhibited significantly slower withdrawal latency relative to each of the PG groups (**Figure 1B**).

Additionally, the Adult-Exposed Heroin rats exhibited significantly slower withdrawal latencies compared with those from the PG groups as well as the Adolescent-Exposed Heroin group, on Days 5 and 10 (**Figure 1B**). A follow up two-way analysis collapsed across Age confirmed main effects of Day (F (2, 60) = 146.89, p < 0.0001), Drug (F (1, 30) = 130.51, p < 0.0001), and an interaction of Day with Drug (F (2, 60) = 141.71, p < 0.0001). The post-hoc confirmed that the PG groups differed from the Heroin groups on Day 1, 5, and 10, and that Day1 differed from Days 5 and 10 within the Heroin groups.

#### Baseline thermal nociception

The three-factor analysis of the first baseline nociception test performed at 12 weeks (PND 85; Adolescent-Exposed groups; **Figure 2A**) or 19 weeks (PND 134; Adult-Exposed groups; **Figure 2B**) of age confirmed only a main effect of Temperature (F (3, 84) = 168.45, p < 0.0001) on withdrawal latency. The post-hoc analysis confirmed that latencies all temperatures differed significantly from one another.

**Figure 2.**
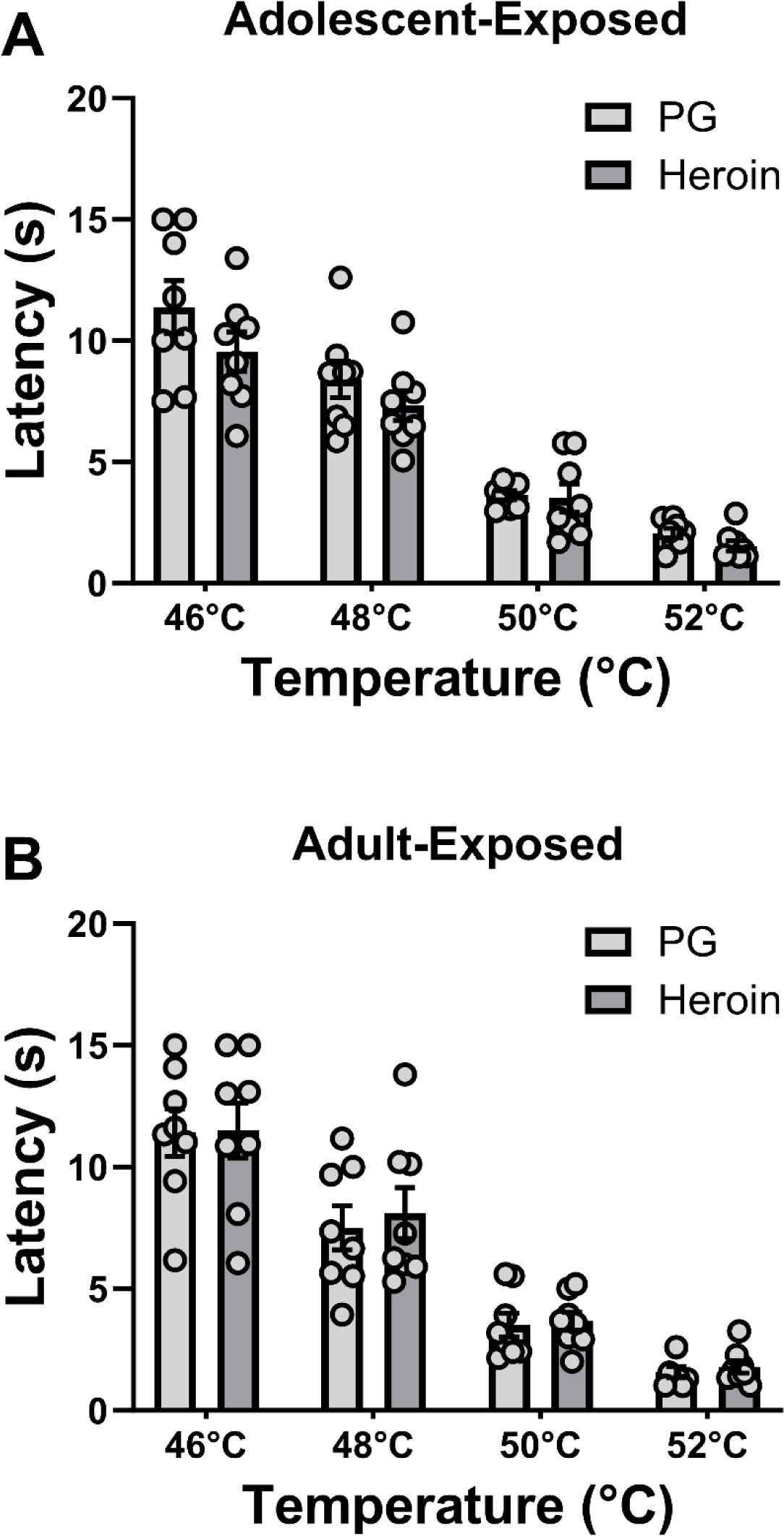
Mean (±SEM) baseline tail-withdrawal latencies for rats repeatedly exposed to vapor during adolescence (*N*=16; 8 PG, 8 Heroin) or adulthood (*N*=16; 8 PG, 8 Heroin) across four temperatures (46°C, 48°C, 50°C, and 52°C). Withdrawal latencies for the Adolescent-Exposed (**A**) and Adult-Exposed (**B**) groups were recorded at 12 and 19 weeks of age, respectively.

A two-factor analysis was performed on baseline nociception data recorded at a second timepoint from the Adolescent-Exposed and Adult-Exposed groups at 17 weeks (PND 121) and 24 weeks (PND 170) of age, respectively. The analysis confirmed only a main effect of Age (F (1, 28) = 4.40, p < 0.05); the Adult-Exposed groups exhibited significantly longer tail-withdrawal latencies compared with the Adolescent-Exposed groups (**Figure 3A**).

**Figure 3.**
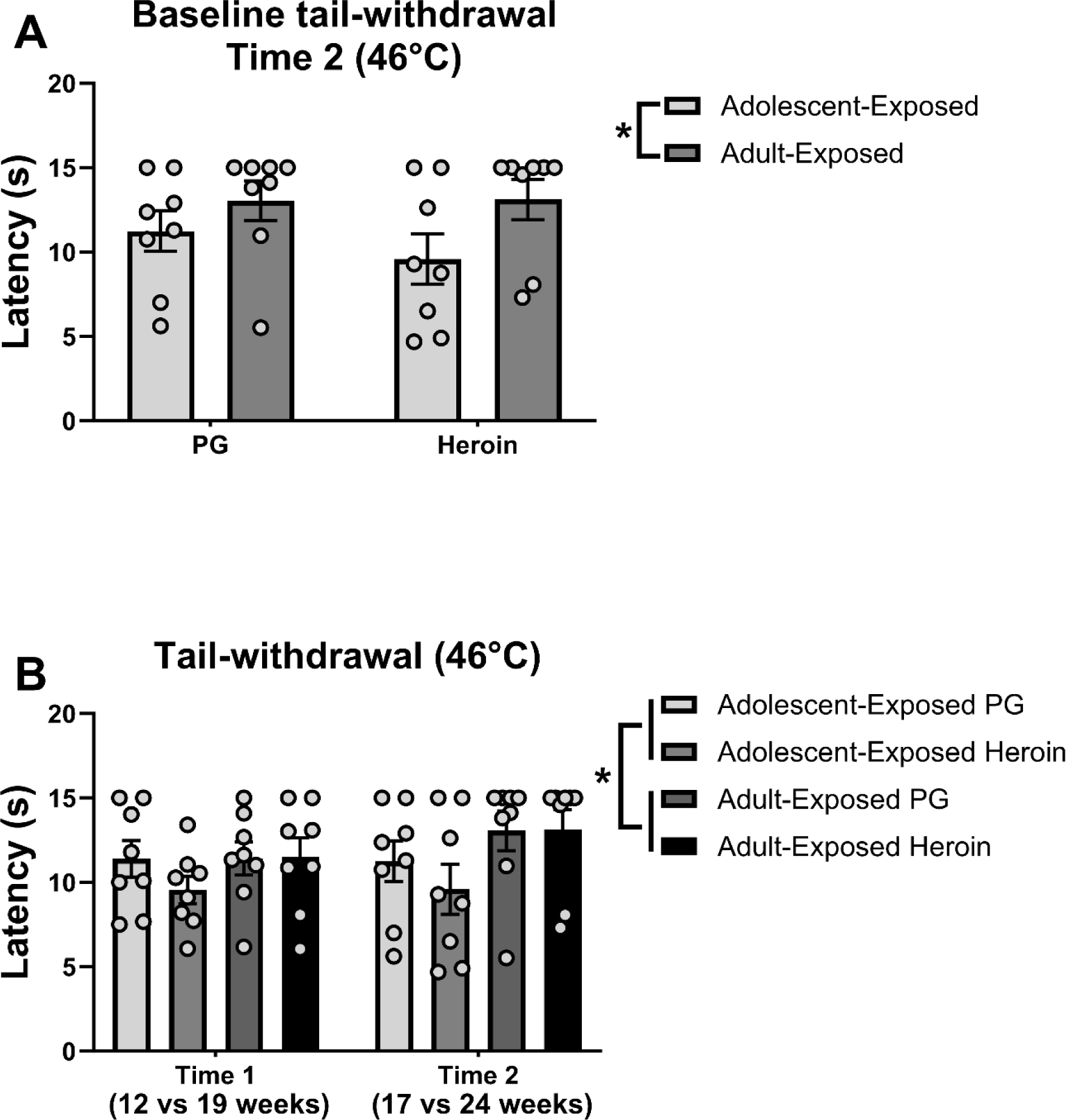
Mean (±SEM) tail-withdrawal latencies during the baseline recording at 17 and 24 weeks of age for the Adolescent-Exposed (*N*=16; 8 PG, 8 Heroin) and Adult-Exposed (*N*=16; 8 PG, 8 Heroin) groups (**A**), respectively. First and second baseline tail-withdrawal measurements taken at 46°C by Treatment Group (**B**). A significant difference between the Adolescent-Exposed and Adult-Exposed groups is indicated with a *.

To determine the effects of Time of assessment, a separate three-way analysis that included both the PND 85 / PND 134 (46 °C only) and PND 121 / PND 170 nociception assessment timepoints was performed. Only a main effect of Age (F (1, 28) = 4.56, p < 0.05) was confirmed, and the withdrawal latencies for the Adult-Exposed groups were significantly longer compared with those from the Adolescent-Exposed groups (**Figure 3B**).

#### Naloxone challenge

An initial three-way analysis, with factors of Treatment condition (pre- and post-naloxone), Age, and Vapor Condition (initial repeated PG/heroin vapor), was performed on tail withdrawal latencies recorded before and after an injection of naloxone (0.3 mg/kg, i.p.). The analysis confirmed significant main effects of Treatment condition (F (1, 28) = 30.81, p < 0.0001) and the interactions of Treatment condition with Age (F (1, 28) = 5.94, p < 0.05) and with Drug (F (1, 28) = 5.20, p < 0.05). The Sidak’s multiple comparison test confirmed that naloxone significantly reduced latencies in all but the rats exposed to heroin during adolescence (**Figure 4A**).

**Figure 4.**
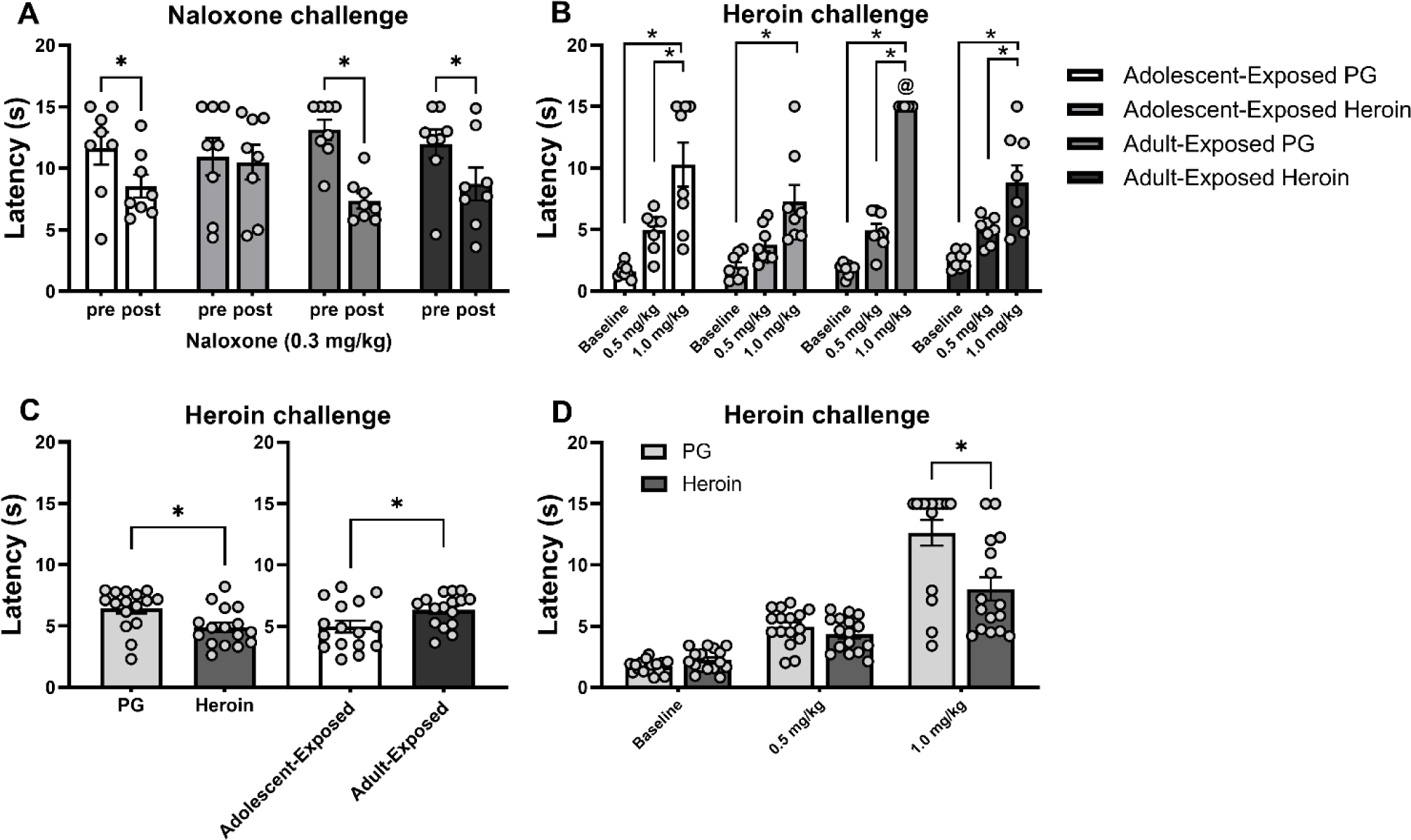
Naloxone and heroin challenges performed on Adolescent-Exposed (*N*=16; 8 PG, 8 Heroin) and Adult-Exposed (*N*=16; 8 PG, 8 Heroin) rats. Tail-withdrawal assessment in (**A**) 46°C water before and after injection of naloxone 0.3 mg/kg i.p. and in (**B-D**) 52°C water before (Baseline) and after injections of heroin 0.5 and 1.0 mg/kg s.c. A significant difference between groups or conditions is indicated with *; and a significant difference compared with all other groups within the same dose with @.

#### Heroin challenge

The initial three-factor analysis of nociception before and after heroin injections confirmed main effects of Treatment condition (F (2, 56) = 117.73, p < 0.0001), Age (F (1, 28) = 6.67, p < 0.05), and of Vapor exposure (F (1, 28) = 8.69, p < 0.01) on withdrawal latencies (**Figure 4B**). Additionally, interactions of Treatment condition with Age (F (2, 56) = 3.83, p < 0.05) and of Vapor exposure with Treatment condition (F (2, 56) = 11.78, p < 0.0001) were confirmed. The marginal means post-hoc analyses further confirmed that all Treatment conditions differed from one another. Additionally, the post-hoc analyses confirmed that the Adolescent-Exposed groups exhibited shorter latencies compared with the Adult-Exposed groups (**Figure 4C-right**) and that latencies of animals in the Heroin groups were shorter than those from animals in the PG groups (**Figure 4C-left**). The Tukey test confirmed no differences in baseline latency between any of the groups nor any differences between baseline and the 0.5 mg/kg dose Treatment conditions within any groups. However, latencies were significantly slower relative to baseline after the 1.0 mg/kg dose within all four groups (**Figure 4B**). Additionally, the Adult-Exposed PG animals showed significantly slower latencies compared with all other groups, after the 1.0 mg/kg heroin injection Treatment condition (**Figure 4B**). The latencies after the 0.5 and 1.0 mg/kg doses significantly differed from each other within all groups except the Adolescent-Exposed Heroin rats (**Figure 4B**). The interactions were further assessed by two-way analyses of each of the two interacting factors collapsed across the third. When Treatment condition and Vapor exposure were considered collapsed across Age, significant effects of Treatment condition (F (2, 60) = 103.70, p < 0.0001), Vapor exposure (F (1, 30) = 7.46, p < 0.05), and the interaction of Treatment with Vapor (F (2, 60) = 10.38, p = 0.0001) were again confirmed. The post-hoc confirmed that the PG and Heroin groups significantly differed but only at the 1.0 mg/kg dose (**Figure 4D**). When the Treatment and Age factors were considered collapsed across previous Vapor exposure, the main effects of Treatment (F (2, 60) = 84.07, p < 0.0001) and Age (F (1, 30) = 5.42, p < 0.05) were once again confirmed; however, the follow-up analysis did not confirm an interaction of factors.

### Anxiety-like behavior

Analysis of *<u>open arm time</u>* in the EPM confirmed main effects of Vapor exposure (F (1, 28) = 5.32, p < 0.05) and Age (F (1, 28) = 9.65, p < 0.01). Rats repeatedly exposed to heroin vapor spent significantly more time in the open arms of the elevated plus-maze compared with those exposed to PG vapor (**Figure 5A**), and rats in the Adolescent groups spent more time in the open arms compared with rats in the Adult groups. The analysis of *<u>total distance traveled</u>* in the EPM apparatus confirmed only a main effect of Age (F (1, 28) = 7.84, p < 0.01); rats in the Adolescent groups traveled significantly more distance compared with rats in the Adult groups, irrespective of Drug vapor treatment (**Figure 5B**).

**Figure 5.**
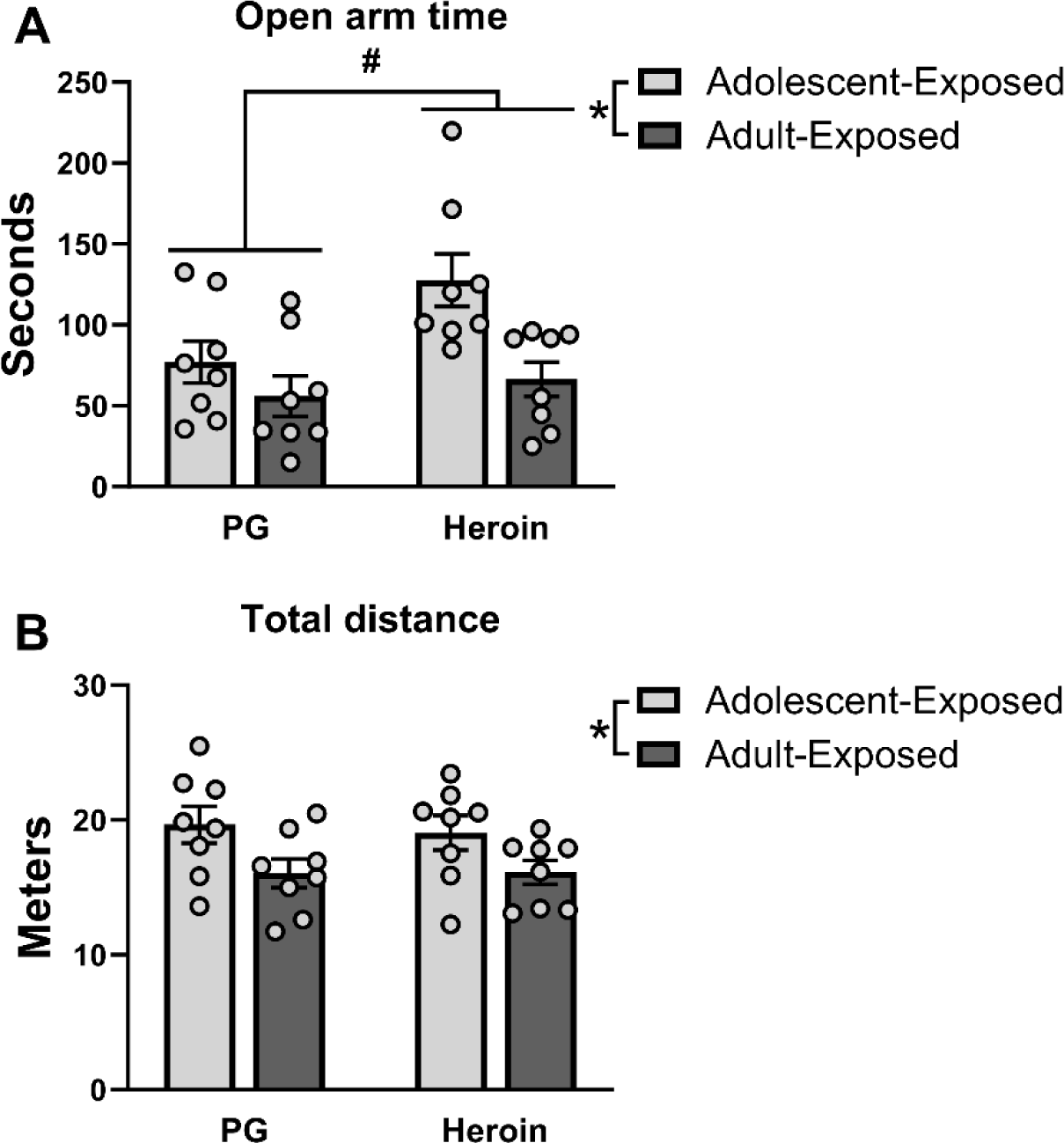
Mean (±SEM) time spent in the open arms (**A**) and total distance traveled (**B**) in the elevated plus-maze for rats repeatedly exposed to PG or heroin vapor during adolescence (*N*=16; 8 PG, 8 Heroin) or adulthood (*N*=16; 8 PG, 8 Heroin). A significant difference between PG and Heroin groups is indicated with #, and a significant difference between Adolescent-Exposed and Adult-Exposed groups with *.

### Spatial learning

The three-factor analysis of *<u>Latency to escape</u>* during the Acquisition Phase in the Barnes maze confirmed a main effect of Day (F (2, 56) = 39.10, p <0.0001) and the marginal means post-hoc further confirmed that latencies for each day significantly differed from those from every other day. The analysis also confirmed an interaction of Age with Day (F (2, 56) = 9.35, p < 0.001). The follow-up two-way considering Age and Day across Vapor exposure again confirmed a main effect of Day (2, 60) = 40.71, p < 0.0001) and the interaction of Age with Day (F (2, 60) = 9.73, p < 0.001). The post-hoc confirmed that the Adult-Exposed animals had significantly longer latencies to escape compared with the Adolescent-Exposed animals on Day 1 (**Figure 6A**).

**Figure 6.**
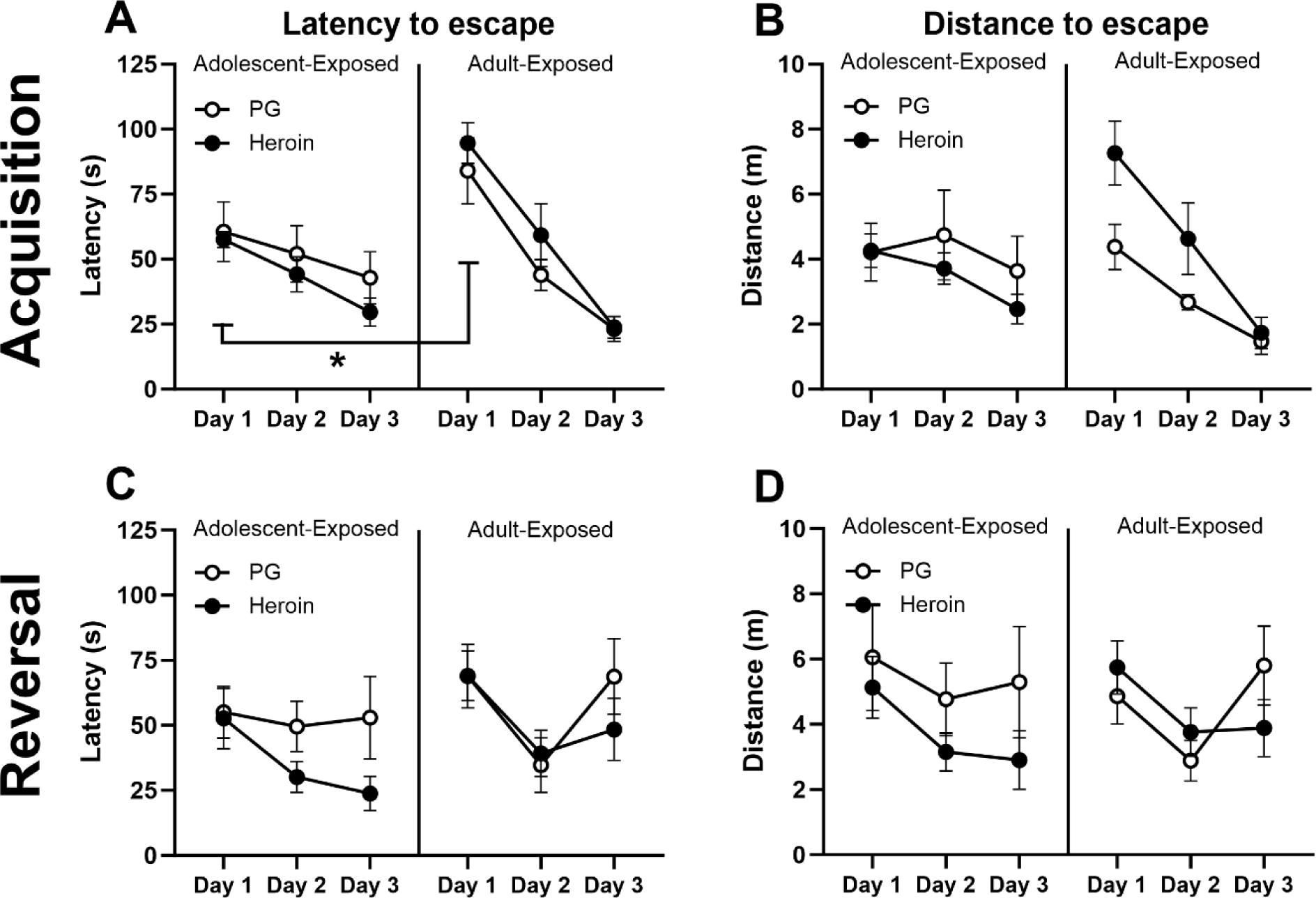
Mean (±SEM) escape latency (**A, C**) and distance traveled to escape (**B, D**) during the acquisition (**A, B**) and reversal (**C, D**) phase of the Barnes maze test. Escape latency and distance traveled to escape was measured in rats repeatedly exposed to vapor during adolescence (*N*=16; 8 PG, 8 Heroin) or adulthood (*N*=16; 8 PG, 8 Heroin). A significant difference between Adolescent-Exposed and Adult-Exposed groups is indicated with *.

The three-factor analysis of the *<u>Distance traveled</u>* to escape during the *Acquisition Phase* confirmed a main effect of Day (F (2, 56) = 15.67, p < 0.0001) and interactions of Age with Day (F (2, 56) = 5.15, p < 0.01) and of Age with Vapor exposure (F (1, 28) = 4.54, p < 0.05). The post-hoc test confirmed that distance traveled was significantly lower on Day 3 compared with Days 1 and 2. A two-way follow-up analysis considering Age and Day across Vapor exposure once again confirmed a main effect of Day (F (2, 60) = 15.49, p < 0.0001) and the interaction of Age with Day (F (2, 60) = 5.09, p < 0.01). The post-hoc did not confirm differences between Age groups on any specific Day (p = 0.058 on Day 1; **Figure 6B**). When Age and Vapor exposure were considered collapsed across Day, the two-way analysis confirmed the interaction of factors (F (1, 28) = 4.54, p < 0.05), however no differences between the groups were confirmed by the post-hoc analysis.

The three-factor analysis of *<u>Latency to escape</u>* during the *Reversal Phase* **(Figure 6C)** confirmed a main effect of Day (F (2, 56) = 6.97, p < 0.01). The post-hoc test confirmed that Latency to escape was significantly reduced on Day 2 compared with Day 1. Similarly, the three-factor analysis of the *<u>Distance traveled</u>* to escape during the *Reversal Phase* **(Figure 6D)** confirmed only a main effect of Day (F (2, 56) = 5.28, p < 0.01). The follow-up post-hoc test for this analysis confirmed that Distance traveled to escape was significantly decreased on Day 2 compared with that of Day 1.

## DISCUSSION

The present study confirmed age-related differences in the development of tolerance during repeated inhalation of heroin vapor and showed that age of exposure can determine the effect of heroin much later. While anti-nociceptive tolerance was observed in both adolescent and adult rats, the adolescent animals exhibited more complete tolerance to heroin (i.e., withdrawal latencies statistically indistinguishable from those of their PG vapor control group) compared with the adults. The effects of repeated heroin vapor inhalation on nociception lasted long past the initial exposure, as evidenced by a lack of effect of naloxone injection in the Adolescent-Exposed heroin inhalation group, and blunted heroin anti-nociception in both Adolescent-Exposed and Adult-Exposed heroin inhalation groups. There was also a long-lasting decrease in anxiety-like behavior associated with repeated heroin inhalation, since both the Adolescent-Exposed and Adult-Exposed groups showed an increase in the amount of time spent in the open arms of the elevated plus maze (EPM). There were also age-related differences confirmed on the first day of the Barnes maze assessment. Importantly, these age-related differences were confirmed throughout the study when all of the animals were in the nominal adult stage. This included when nociception was probed at 17 versus 24 weeks of age, when groups were challenged with heroin (21 vs 28 weeks of age), and when anxiety-like behavior was evaluated (13 vs 20 weeks of age).

The nociceptive tolerance observed in this study after vapor inhalation of heroin is consistent with previous work showing that repeated subcutaneous injections of morphine during adolescence induces tolerance at a faster rate compared with treatment during adulthood (Nozaki et al. 1975) and that opioid anti-nociception is blunted in adulthood in animals exposed to oxycodone during adolescence by way of intravenous self-administration (Zhang et al. 2016). The present study extends these previous findings in male rats to female rats. Importantly, the present study also shows that nociceptive effects of repeated opioids, which were confirmed out to ∼16 weeks after the initial heroin vapor exposure in the Adolescent and Adult groups, are persistent. This is consistent with our previous work showing that repeated exposure to heroin vapor during adolescence increases baseline sensitivity to thermal stimuli and reduces the acute analgesic effects of subcutaneous heroin injections, in adulthood (Gutierrez et al. 2022a). This study also extends prior results by showing that while the blunting of heroin-induced anti-nociception was present in both Adolescent-Exposed and Adult-Exposed groups, the *hyperalgesic* effects of the opioid antagonist naloxone were absent *only* in the Adolescent-Exposed group. This may be related to the more-complete tolerance induced during the exposure phase but future study of different exposure conditions to produce similar levels of tolerance across ages would be required to fully explore this interpretation.

Prior studies of adolescent opioid exposure on the expression of anxiety-like behavior later in life report mixed results. For example, no effects on anxiety-like behavior were found in the light-dark and open-field tests in mice after repeated injections (i.p.) of morphine (Lutz et al. 2013), in the open-field and marble burying tests in male mice after chronic delivery of oxycodone via osmotic minipumps (Sanchez et al. 2016), or in the EPM after repeated injections (i.p. and s.c.) of morphine in male rats (Schwarz and Bilbo 2013). The latter study did find a decrease in EPM open arm time when the same dosing regimen was administered to male rats in during adulthood (Schwarz and Bilbo 2013). However, a more recent study found that repeated s.c. injections of morphine during adolescence produced an *increase* in open arm time in the EPM in adulthood in male rats (Khani et al. 2022a). In the present study, we found that repeated exposure to heroin vapor reduced anxiety-like behavior, as measured using the EPM test. Although there was not any confirmed statistical difference associated with age of heroin inhalation, qualitatively the Adolescent-Exposed group exhibited an anxiolytic effect more consistently, and of greater magnitude, than did the Adult-Exposed group. The reason for the discrepancies in prior findings is not clear, but potential causes include methodological differences, such as the stage of adolescence during which exposure occurred, dosing or testing procedures, or factors such as ambient lighting or the time of day when the behavioral testing was carried out. Our finding was particularly interesting given that we previously reported an *increase* in anxiety-like behavior following repeated adolescent exposure to heroin vapor (Gutierrez et al. 2022a). Our studies were identical on several factors such as the dose, route of administration, age of treatment, strain of rat, etc., which limits the possible contributing factors. One potential difference between our two studies was that dosing was performed in a well-lit room, and testing occurred in different rooms, for the first study, while they occurred within the same location in the present study with dosing performed in the dark under red light. Further determination of the circumstances that produce increased versus decreased anxiety-like behavior, while important, is outside of the scope of this study, which is focused on contrasting the age of exposure. In general, the evidence reported here supports the conclusion that repeated adolescent and adult opioid exposure can each have a lasting impact on the expression of anxiety-like behavior.

Chronic opioid use has been associated with deficits in learning and memory (Baldacchino et al. 2018; Mitrovic et al. 2011) even after long periods of abstinence (Schmidt et al. 2017), in adult humans. However, there is minimal information about the long-term effects of repeated *adolescent* opioid exposure on learning and memory. This is also true for animal studies, where the focus has primarily been on the acute effects, effects observed during chronic treatment, or those seen shortly after drug discontinuation or upon reaching early adulthood. Additionally, the bulk of prior studies have focused on effects in male rodents. In the present study, we assessed spatial navigation in adult female rats exposed to heroin or PG vehicle during adolescence or adulthood using a Barnes maze and found no differences in performance associated with the repeated heroin inhalation. This lack of effect contrasts with a recent study which reported that repeated injections (s.c.) of morphine during adolescence significantly impacted spatial learning and memory during adulthood in rats (Khani et al. 2022b). In this previous study, the authors used a Morris water maze and reported effects of morphine in acquisition, as well as in the memory retention probe. Differences in methodology could account for some of the differences seen between these two studies, including in the motivation to perform in each specific behavioral test and the sex of the animal subjects. One minor limitation of the current study was the lack of a probe for memory retention following the acquisition and reversal phases since some previous work has shown that repeated exposure to opioids can impact memory retention while sparing acquisition performance (Brolin et al. 2018); it would be of interest to assess retention in future studies. Finally, the Barnes maze is interpreted as a test of learning and memory but depends on negative reinforcement. The escape into the goal box is reinforced by the removal of the anxiogenic properties of the large open surface of the maze. Thus, given the reduction of anxiety caused by repeated heroin vapor, as assessed with the EPM, it is possible that any group differences in learning or memory may have been obscured by the differential value of the negative reward.

In conclusion, the data reported in this paper provide evidence that both adolescent and adult heroin exposure by way of inhalation can result in long-term alterations in nociceptive sensitivity and can decrease the expression of anxiety-like behavior. It should be noted that age-specific effects, although minor, were confirmed in this study and included an insensitivity to naloxone-induced hyperalgesia and a lack of a difference in tail-withdrawal latency between the two injected heroin doses in the Adolescent-Exposed Heroin group. Furthermore, while there was no statistical difference in EPM open arm time between Adolescent- and Adult-Exposed Heroin groups, the degree to which heroin increased open arm time in the Adolescent-Exposed group was greater. While these observations may warrant future investigations on the age-specific effects of opioid exposure, the major finding of this study is that repeated heroin vapor inhalation can produce long-lasting effects in both adolescent- and adult-exposed rodents.

## Acknowledgements

We would like to thank Ana D.A. Gahng for her assistance in conducting portions of the behavioral experiments.

